# Targeted stimulation of an orbitofrontal network disrupts decisions based on inferred, not experienced outcomes

**DOI:** 10.1101/2020.04.24.059808

**Authors:** Fang Wang, James D. Howard, Joel L. Voss, Geoffrey Schoenbaum, Thorsten Kahnt

## Abstract

When direct experience is unavailable, animals and humans can imagine or infer the future to guide decisions. Behavior based on direct experience versus inference may recruit distinct but overlapping brain circuits. In rodents, the orbitofrontal cortex (OFC) contains neural signatures of inferred outcomes, and OFC is necessary for behavior that requires inference but not for responding driven by direct experience. In humans, OFC activity is also correlated with inferred outcomes, but it is unclear whether OFC activity is required for inference-based behavior. To test this, we used non-invasive network-based continuous theta burst stimulation (cTBS) to target lateral OFC networks in the context of a sensory preconditioning task that was designed to isolate inference-based behavior from responding that can be based on direct experience alone. We show that relative to sham, cTBS targeting this network impairs reward-related behavior in conditions in which outcome expectations have to be mentally inferred. In contrast, OFC-targeted stimulation does not impair behavior that can be based on previously experienced stimulus-outcome associations. These findings suggest that activity in the targeted OFC network supports decision making when outcomes have to be mentally simulated, providing converging cross-species evidence for a critical role of OFC in model-based but not model-free control of behavior.

## INTRODUCTION

Many decisions are made based on expectations about their likely outcomes. Such expectations can reflect what we have experienced in the past, for instance, when ordering your favorite dish at a familiar restaurant. For many other decisions in life, such as deciding to try out a new restaurant or enrolling in a PhD program, direct experience is lacking, and outcome expectations need to be mentally simulated or inferred.

Expectations arising from these two different origins, which may compete for control over behavior (Daw et al., 2005; Lee et al., 2014), are thought to recruit distinct but overlapping brain circuits (Balleine and Dickinson, 1998; Daw et al., 2011; O’Doherty et al., 2017). Whereas much research has focused on behavior that is based on direct experience (Schultz, 1998; Tricomi et al., 2009; Wunderlich et al., 2012), less is known about the neural representations that support behavior based on inferred outcomes in humans.

Work across animal species suggests that the orbitofrontal cortex (OFC), together with the hippocampus, is particularly important for behavior based on inference (Rudebeck and Murray, 2014; Wikenheiser and Schoenbaum, 2016). For instance, in tasks that require mental simulation, neural activity in the rodent OFC represents inferred outcomes in almost the same way as it signals directly experienced outcomes (Takahashi et al., 2013; Sadacca et al., 2018). Interestingly, however, the rat OFC is not required for behavior based on directly experienced outcomes, but it is only necessary when responding requires inference (Jones et al., 2012; Takahashi et al., 2013). This suggests that rodent OFC is selectively required for the simulation of outcomes. Recent work in humans has shown similar neural correlates of inferred outcomes in the OFC (Barron et al., 2013; Wang et al., 2020), but whether human OFC networks are required for behavior based on such inferred outcomes is unclear.

Causal studies on human OFC function have traditionally been limited to naturally-occurring lesions (Reber et al., 2017; Vaidya et al., 2019). However, we have recently developed a novel network-based transcranial magnetic stimulation (TMS) approach to non-invasively target activity in human OFC networks (Howard et al., 2020). Similar to previous work targeting the hippocampal network (Wang et al., 2014), this approach uses resting-state functional magnetic resonance imaging (rsfMRI) to individually define stimulation coordinates in the lateral prefrontal cortex (LPFC) that are part of the central/lateral OFC network (Kahnt et al., 2012; Zald et al., 2014). We have recently shown that this targeted TMS protocol selectively affects connectivity in lateral OFC networks, in parallel with disrupting OFC-dependent behavior (Howard et al., 2020).

In the current study (**Fig. 1A**), we applied this novel OFC-targeted brain stimulation approach in the context of a sensory preconditioning task that was designed to isolate inference-based behavior from responding that can be based on direct experience (Jones et al., 2012; Wimmer and Shohamy, 2012; Wang et al., 2020). This task consists of three phases (**Fig. 1B**): First, during preconditioning, pairs of sensory cues are repeatedly presented (A⟶B, C⟶D). Next, during conditioning, the second cue of each pair is associated with reward and no reward, respectively (B⟶reward, D⟶no reward). During the final probe test, reward-related responding to each cue (A, B, C, and D) is probed under extinction conditions. Reward-related responses to cue A indicate that subjects step through the associations A⟶B and B⟶reward to infer A⟶reward. In contrast, such responses to cue B do not require inference because direct experience with the cue-outcome pairing is available. We predicted that disrupting OFC network activity with OFC-targeted TMS will impair inference-based behavior (responding to cue A), but not behaviors that can be based entirely on direct experience alone (responding to cue B).

**Figure 1.**
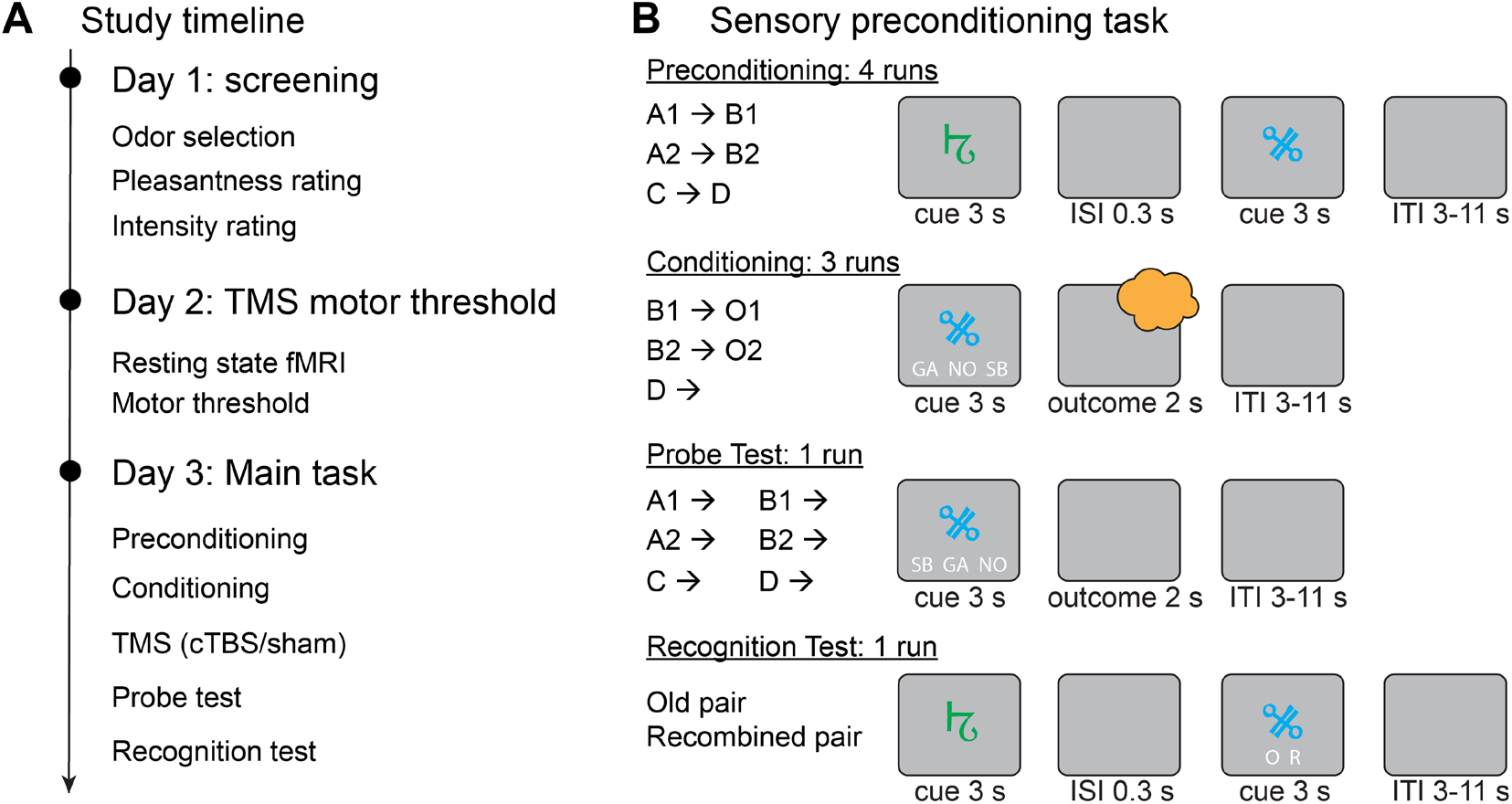
Experimental design and sensory preconditioning task. **A.** Experimental timeline. **B.** Participants learned cue pairs during preconditioning (A⟶B, C⟶D). During conditioning, they learned associations between the second cue in each pair and one of two food odors (O1 or O2) or odorless air (B ⟶ odor reward, D ⟶ odorless air). During the probe test, participants were asked to make outcome predictions to all cues, but no outcomes were delivered. Finally, subjects completed a recognition task testing for memory of cue-cue associations.

## MATERIALS AND METHODS

### Subjects

In total, 71 healthy adults participated in a screening session. Of these, 52 passed screening, were randomly assigned to the sham (SHAM: n=25, 13 female) or stimulation group (STIM: n=27, 15 female), and participated in the experiment. All participants provided written informed consent to participate and were compensated with $20 per hour for behavioral testing and $40 per hour for TMS and MRI scanning. The study protocol was approved by the Northwestern University Institutional Review Board. One participant in the STIM group withdrew during the experiment. Data from four participants (two per group) were excluded from all analyses because their performance in the last run of conditioning was not significantly above chance (p>0.05, binomial test). This left a total of 47 participants (SHAM: n=23, 12 female, mean age=25.24 years ± 0.86 s.e.m; STIM: n=24, 13 female, age=25.30 years ± 0.70) from whom data was analyzed. Of those, data from four participants (one SHAM, three STIM) from the recognition memory test of the experiment were not recorded due to technical problems.

### Stimuli and Odor Delivery

Visual cues consisted of 14 abstract symbols, and 12 of them were randomly grouped into six pairs for each participant. Two pairs served as A1–B1 pairs, two served as A2–B2 pairs, and two served as C–D pairs. The two remaining symbols were used to form two catch-trial pairs (E–E) in which the same symbols were presented twice in a row (i.e., E1–E1, E2–E2). The two symbols constituting a pair were presented in different colors (e.g., first symbol blue, second symbol green; counterbalanced across participants).

As in our previous studies, the current experiment used food odors as biologically relevant reward in hungry participants (Howard et al., 2015; Howard and Kahnt, 2017, 2018; Suarez et al., 2019; Howard et al., 2020). Eight food odors (four sweet: strawberry, caramel, gingerbread, and yellow cake; four savory: potato chip, pot roast, garlic, and pizza) were provided by Kerry (Melrose Park, IL) and International Flavors and Fragrances (New York, NY). Odors were delivered to participants’ nose using a custom-built and computer-controlled olfactometer (Howard et al., 2020; Wang et al., 2020). The olfactometer was equipped with two independent mass flow controllers (Alicat, Tucson, AZ), which allow dilution of any given odorant with odorless air. Odorless air was delivered constantly during the experiment and odorized air was mixed into the airstream at specific time points. The overall flow rate was kept constant at 3.2 L/min throughout the task, such that odor deliver did not involve a change in overall airflow or any noticeable change in somatosensory stimulation.

### Experimental Task and Design

The study was conducted over three days (**Fig. 1A**) and included (1) a screening session, (2) a MRI and TMS motor threshold session, and (3) a main task session. The MRI and TMS motor threshold session was conducted on average 18 days (s.e.m.=4.16) after the screening session. And the average delay between motor threshold and main task sessions was 4 days (s.e.m.=0.94). Participants were instructed to arrive in a hungry state (fast for at least 4 hours) for the screening and main task sessions.

#### Screening Session

After informed consent and screening for eligibility, participants’ rated the pleasantness of eight food odors. In each trial, participants were presented with one of the eight food odors for 2 seconds and were instructed to make a medium sized sniff. They then rated the pleasantness of the delivered odor on a scale from “Most disliked sensation” to “Most liked sensation”. Each food odor was presented 3 times in randomized order and ratings were averaged. We then selected one sweet and one savory odor that were both rated as pleasant (i.e. pleasantness above neutral) and as closely matched as possible. The two selected odors were then used as reward for that individual participant in the main task session. If no such two odors were found, participants were excluded from further participation in the study. Next, participants rated the intensity and pleasantness of the two selected odors as well as odorless air. The scale of the intensity rating was from “Undetectable” to “Strongest sensation imaginable”.

#### MRI and TMS Motor Threshold Session

We acquired a T1-weighted structural MRI scan for the purpose of TMS neuronavigaton and an 8.5 minutes rsfMRI scan for individually defining OFC-targeted stimulation coordinates (see below). We then measured resting motor threshold (RMT) by delivering single TMS pulses over left motor cortex. RMT was defined as the minimum percentage of stimulator output necessary to evoke 5 visible thumb movements in 10 stimulations.

#### Main Task Session

The main task session consisted of preconditioning, conditioning, TMS, probe test, and a cue-cue recognition memory test (**Fig. 1B**). In four preconditioning runs, participants were instructed to learn the associations between the two cues in each pair (A⟶B [A1⟶B1, A2⟶B2], C⟶D [C1⟶D1, C2⟶D2], E⟶E [E1⟶E1, E2⟶E2]). The cues in a given pair were presented one after another for 3 seconds each, separated by an inter-stimulus-interval (ISI) of 300 ms. A fixation cross appeared between trials for a variable inter-trial-interval (ITI) between 3 and 11 seconds. To ensure attention to the cue pairs, participants were instructed to memorize the cue pairs, press a button if the second cue was different from the first cue, and withhold a response if the two cues were identical. To facilitate learning, in the first two runs of preconditioning, each cue pair was repeated three times in a row. In the remaining preconditioning runs, the order of cue pairs was randomized across trials.

Next, participants performed three runs of conditioning, during which the second cue of each pair (cues B [B1, B2] and D [D1, D2]) was presented individually for 3 seconds. Participants were instructed to indicate by button press which outcome (e.g. strawberry odor [SB], garlic odor [GA], or no odor [NO]) they expected following the cue. If they expected strawberry, they were asked to select “SB”; if they expected garlic, they were asked to select “GA”; If they expected no odor, they were asked to select “NO”. Participants made their prediction by pressing a button with the index, middle or ring fingers of their right hand corresponding to the positions of “SB”, “GA” and “NO” on the screen. The positions of the abbreviated names were randomized across trials. Irrespective of their selection, the outcome was presented for 2 seconds immediately after the cue. However, “too slow” was displayed if participants failed to respond within 3 seconds. Each cue-outcome association was repeated four times in each run in pseudorandomized order.

After the conditioning phase, participants received OFC-targeted cTBS (see below). The probe test followed immediately after the stimulation. In each trial of the probe test, cue A (A1, A2), B (B1, B2), C (C1, C2), or D (D1, D2) was presented individually under extinction conditions (odorless air was delivered throughout) to prevent further learning. Each cue was presented four times in pseudorandomized order. Participants were instructed to predict the outcome after each cue, as they did during the conditioning phase. They were further instructed to use the cue-cue associations learned in the first phase to infer the outcomes associated with the preconditioned cues (Wang et al., 2020). The durations of cue presentation and the ITI were the same as during the conditioning phase.

Following the probe test, participants were tested for their memory of the cue-cue associations in a recognition task. On each trial, participants were presented with either an original cue pair or with a newly recombined pair (i.e., consisting of cues belonging to different pairs). Pairs were presented sequentially as during preconditioning, and participants were asked to indicate using a button press whether a pair was old (O) or recombined (R) after the second cue was presented.

### MRI Data Acquisition

MRI data were acquired at the Northwestern University Center for Translational Imaging (CTI) using a Siemens 3T PRISMA system equipped with a 64-channel head coil. rsfMRI scans were acquired with an echoplanar imaging (EPI) sequence with the following parameters: repetition time (TR), 2 s; echo time (TE), 22 ms; flip angle, 90°; slice thickness, 2 mm, no gap; number of slices, 58; interleaved slice acquisition order; matrix size, 104 × 96 voxels; field of view, 208 mm × 192 mm; multiband factor, 2. To minimize susceptibility artifacts in the OFC, the acquisition plane was tilted approximately 25° from the anterior commissure (AC)–posterior commissure (PC) line. The rsfMRI scan consisted of 250 EPI volumes covering all but the most dorsal portion of the parietal lobes. In addition, a 3D 1 mm isotropic T1-weighted structural scan was also collected (TR, 2300 ms; TE, 2.94 ms; flip angle, 9°; field of view, 176 mm × 256 mm × 256 mm)

### fMRI Data Preprocessing

Preprocessing of functional images was performed using Statistical Parametric Mapping (SPM12, https://www.fil.ion.ucl.ac.uk/spm/). To correct for head motion during scanning, all rsfMRI images were aligned to the first acquired image. The mean realigned images were then co-registered to the anatomical image, and the resulting registration parameters were applied to all realigned EPI images. Finally, co-registered EPI images were resliced and smoothed with a 6 × 6 × 6 mm Gaussian kernel. To generate forward and inverse deformation fields, the anatomical image was normalized to Montreal Neurological Institute (MNI) space using the 6-tissue probability map provided by SPM12.

### OFC-targeted Transcranial Magnetic Stimulation

We used our previously established network-based OFC-targeted TMS protocol (Howard et al., 2020). TMS was delivered using a MagPro X100 stimulator connected to a MagPro Cool-B65 butterfly coil (MagVenture A/S, Farum, Denmark). We used a cTBS protocol involving a 40 second train of 3-pulse 50 Hz bursts delivered every 200 ms (5 Hz), totaling 600 pulses (Huang et al., 2005). This TMS protocol has inhibitory aftereffects that last for 50-60 minutes over motor cortex (Huang et al., 2005). Stimulation was delivered at an intensity of 80% MT in the STIM group and 5% MT in the SHAM group. As in our previous study (Howard et al., 2020), the target coordinate was defined as a location in the right LPFC that showed maximal functional connectivity with the right OFC seed coordinate (see details below). The orientation of the coil was such that the long axis of the figure-of-eight coil was approximately parallel to the long axis of the middle frontal gyrus. All participants were informed that they may experience muscle twitches in the forehead, eye area, and jaw during stimulation. We delivered two single test pulses to test for tolerability before cTBS was delivered. Immediately after the last pulse of cTBS, the time was noted. All subsequent testing (probe test and recognition memory) took place within 33 ± 1.92 minutes of the end of TMS, and this time did not differ between groups (t=0.24, p=0.814).

### Coordinate selection for OFC-targeted TMS

Stimulation coordinates on the right LPFC were determined for each individual participant based on rsfMRI connectivity with a right central-lateral OFC seed region using a previously described procedure (Howard et al., 2020). Briefly, we first created two spherical masks of 8-mm radius around a LPFC coordinate (x=48, y=38, z=20) and a OFC seed coordinate (x=28, y=38, z=−16) in MNI space, both inclusively masked by the gray matter tissue probability map provided by SPM12 (thresholded at >0.1). These masks were then inverse-normalized to each participant’s native space using the inverse deformation field generated by normalizing the anatomical image. We then estimated a general linear model with the average rsfMRI time series in the OFC mask as the regressor of interest and realignment parameters as regressors of no interest. The voxel in the LPFC mask that had highest functional connectivity with the OFC seed was defined as stimulation coordinate. We used neuronavigation to apply stimulation to this coordinate.

### Statistics

Simple between-group effects were tested using unpaired t-tests. Results from parametric tests were confirmed using permutation tests involving 10,000 random group assignments. Interactions were tested using R (R Core Team, 2018) and the lme4 package (Bates, 2010). Specifically, we performed linear mixed effect analysis on odor pleasantness ratings with group (SHAM vs. STIM) and odor (odor vs. odorless) as independent variables. In addition, to test the interaction between group, cue type, and time on reward predictions during conditioning, we used a generalized linear mixed model with group (SHAM vs. STIM), cue (B vs. D), and time (three runs) as independent variables. Finally, the interaction between group and cue type on reward predictions during the probe test was tested using a generalized linear mixed model with group (SHAM vs. STIM) and cue type (A vs. B) as independent variables. In all analyses, subjects were modeled as random intercept effects. There were no obvious deviations from normality or homoscedasticity based on visual inspection of residual plots. We computed p values by likelihood ratio tests (χ^2^) of the full model including the effect of interest against the reduced model without the effect of interest. Statistical thresholds were set to p<0.05, two-tailed unless indicated otherwise.

## RESULTS

### Odor Ratings and Learning Performance

The experiment took place across three days (**Fig. 1A**). Day 1 and 2 consisted of a screening visit and a MRI (anatomical and rsfMRI) session, respectively. Day 3 involved a sensory preconditioning task and network-based OFC-targeted TMS. On day 3, subjects (SHAM, n=23; STIM, n=24) in both groups arrived fasted (not eaten for 11 ± 4.27 hours; group difference, t(45)=1.00, p=0.321) and with similar levels of hunger (t(45)=1.28, p=0.205). Subjects first learned associations between pairs of abstract visual cues during preconditioning (A⟶B, C⟶D, **Fig. 1B**). Next, they learned that a pleasant food odor followed cue B, whereas cue D was always followed by odorless air (**Fig. 1B**). To measure reward expectations, participants were asked to predict the outcome associated with the presented cue via button press.

Subjects in both groups rated the food odors as significantly more pleasant than the odorless air (SHAM: t(22)=11.62, p=7.38×10^−11^; STIM: t(23)=12.97, p=4.59×10^−12^, **Fig. 2A**), demonstrating that food odors were perceived as rewarding. Importantly, there were no differences in the pleasantness ratings between groups (main effect of group: χ^2^(1)=2.49, p=0.115; group by odor interaction: χ^2^(1)=1.34, p=0.247). During conditioning, the percentage of trials in which participants expected a food odor after cue B increased across time relative to cue D (3-way [group × time × cue] generalized linear mixed model; main effect of cue, χ^2^(1)=1736, p<2.2×10^−16^; main effect of time, χ^2^(2)=0.98, p=0.613; cue by time interaction, χ^2^(2)=254.22, p<2.2×10^−16^; **Fig. 2B**). There were no significant differences between groups in learning across time (main effect of group, χ^2^(1)=0.096, p=0.757; cue by group interaction, χ^2^(1)=3.22, p=0.072; time by group interaction, χ^2^(2)=2.88, p=0.24; cue by time by group interaction, χ^2^(2)=0.36, p=0.834). Most importantly, performance in the last conditioning run did not differ between groups (t(45)=0.0045, p=0.996), demonstrating that subjects in both groups learned the associations between the cues and their associated outcomes equally well.

**Figure 2.**
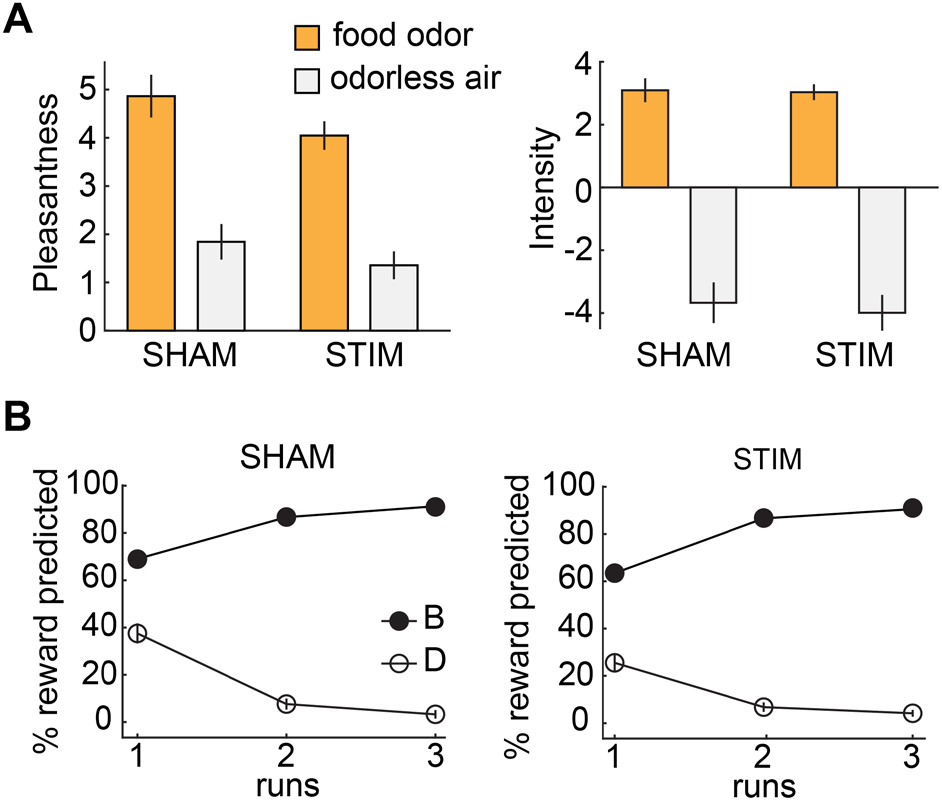
Odor ratings and behavioral performance during conditioning. **A.** Participants rated the pleasantness (left) and intensity (right) of food odors significantly higher than odorless air (p<0.001), but ratings did not differ between groups (p’s>0.14). **B.** The percentage of trials in which an odor reward was expected after cue B increased relative to cue D across time during conditioning, and there were no group differences. Error bars depict SEM (n=23 SHAM, n=24 STIM).

### OFC-targeted cTBS disrupts inference-based responding

After conditioning and immediately prior to the probe test, we applied 40 seconds of cTBS to a site in right lateral prefrontal cortex (LPFC) that was individually selected to have maximal resting-state fMRI connectivity with the central/lateral OFC, following previously established procedures (Howard et al., 2020). Specifically, stimulation was administered in the STIM group at a high intensity that we have previously shown disrupts OFC network activity and adaptive behavior in a reinforcer devaluation task. Stimulation in the SHAM group was administered at a low intensity that was not expected to produce any impact on neural function (Howard et al., 2020).

We hypothesized that targeting the lateral OFC network with cTBS would selectively disrupt reward expectations based on inference but not those based on direct experience. In line with this, we found a significant interaction between cue type and group (χ^2^(1)=4.95, p=0.026), indicating that responses to cues A and B were differentially affected by OFC-targeted cTBS compared to the SHAM group. Indeed, follow-up t-tests showed that this interaction was driven by significantly reduced responses to cue A in the STIM relative to the SHAM group (t(45)=2.40, p=0.020, **Fig. 3A**) whereas there was no group difference in responding to cue B (t(45)=1.18, p=0.245, **Fig. 3B**). These results were confirmed using permutation tests (group difference in responding to A, p=0.012; group difference in responding to B p=0.127). This demonstrates that effects of OFC-targeted cTBS were specific for inference-based responding.

**Figure 3.**
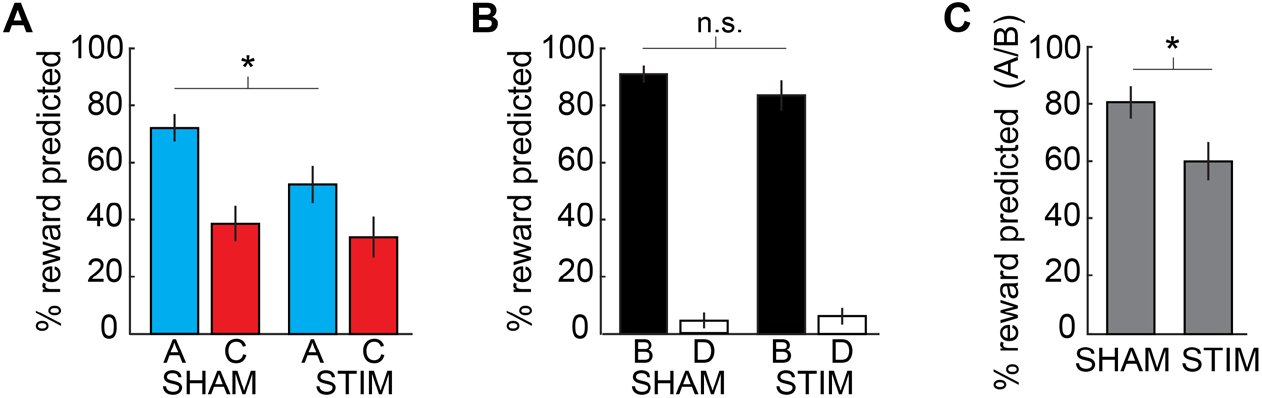
Responses based on inferred outcomes are disrupted by OFC-targeted cTBS. **A.** The percentage of trials in which participants predicted a reward for cue A was significantly larger in the SHAM compared to the STIM group (p=0.020). There was no difference in reward predictions for cue C (p=0.642). **B.** There was no group difference in responding to cue B (p=0.245) or D (p=0.740). **C.** Responses to cue A relative to cue B (A/B) were significantly stronger in the SHAM compared to the STIM group (p=0.024). Error bars depict SEM (n=23 SHAM, n=24 STIM) and * depicts p<0.05.

Reward-related responding to cue A depends not only on the ability to make an inference, but also on knowledge about the reward predicted by cue B, which was acquired through direct experience (B⟶reward). To further examine the effects of OFC-targeted cTBS on inference-based behavior independent of potential effects on direct experience, we normalized responses to cue A by responses to cue B. The resulting ratio (i.e., A/B) reflects the ability to infer outcomes relative to the knowledge about directly experienced cue-reward association. This ratio was significantly smaller in the STIM compared to the SHAM group (t(45)=2.33 p=0.024, **Fig. 3C**). We confirmed the statistical significance of this difference using a permutation test (p=0.013). Taken together, these results demonstrate that OFC-targeted cTBS selectively impairs behavior based on inferred outcomes but does not disrupt behavior that can be based on directly experienced outcomes.

### OFC-targeted cTBS does not disrupt memory for cue-cue associations

Inference also depends on memory of the cue-cue associations learned during preconditioning (Wang et al., 2020). It is therefore possible that the findings reported above reflect a failure of memory rather than inference. Although this is unlikely given that memory of directly experienced cue-reward associations was unimpaired in the STIM group, we measured recognition memory for cue-cue associations after the probe test to rule out this potential explanation. In both groups, recognition memory was significantly above chance (SHAM: t(21)=5.01, p<0.001; STIM: t(20)=2.70, p=0.013), and there was no difference between groups (t(41)=1.34, p=0.188, permutation test, p=0.129 **Figure 4A**). Moreover, as in our previous study (Wang et al., 2020), recognition memory was significantly correlated with inference-based responding (r=0.51, p=0.0005, **Figure 4B**). These correlations were significant within each group (SHAM, r=0.38, p=0.039, one-tailed; STIM, r=0.55, p=0.01) and did not differ between groups (Z=−0.93, p=0.178). Taken together, these findings demonstrate that similar to directly experienced cue-reward associations, OFC-targeted cTBS did not significantly impair memory for cue-cue associations, or how they were used for inference-based behavior.

**Figure 4.**
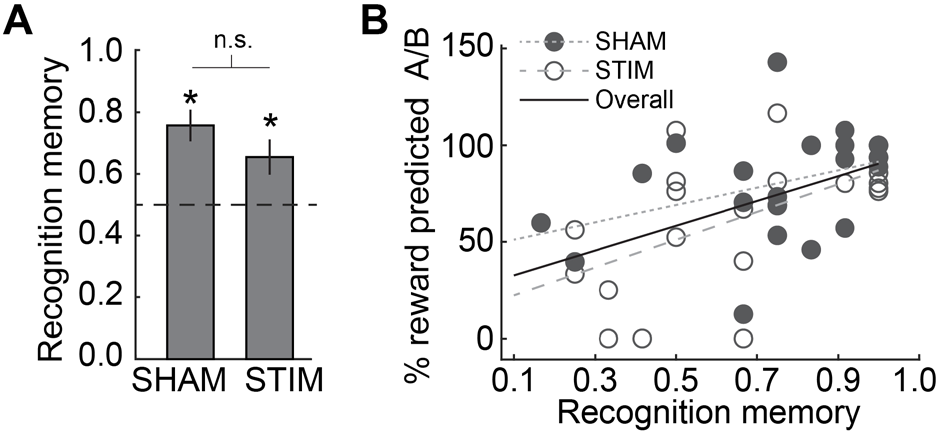
Memory for cue-cue associations and its relation to inference-based behavior is not altered by OFC-targeted cTBS. **A.** Recognition memory for cue-cue pairs does not differ between groups (p=0.188). * depicts p<0.05. **B.** Recognition memory for cue-cue associations was significantly correlated with responding to preconditioned cues (reward prediction responses to A/B) during the probe test (r=0.51, p<0.001, solid circles: SHAM; empty circles: STIM), and this correlation did not differ between groups (Z=−0.93, p=0.178).

## DISCUSSION

The current study shows that targeting the human OFC with network-based cTBS impairs reward-related behaviors when outcome expectations need to be mentally simulated, but not when expectations can be based on direct experience. This closely parallels previous findings from rats (Jones et al., 2012), providing converging cross-species evidence for a critical role of OFC networks in model-based but not model-free behavior.

As such, our findings suggest that the contribution of OFC to decision making may be limited to situations that require model-based planning, and that choices based on direct experience may rely on value computations in other brain areas, such as the amygdala or striatum (Paton et al., 2006; Cox and Witten, 2019). This proposal is seemingly at odds with a large number of studies across different species that has shown neural correlates of both inferred and directly experienced value in OFC (Hare et al., 2009; Schoenbaum et al., 2009; Barron et al., 2013; Stalnaker et al., 2014; Howard et al., 2015; Padoa-Schioppa and Conen, 2017; Suzuki et al., 2017; Klein-Flugge et al., 2019; Lopez-Persem et al., 2020; Wang et al., 2020). Why would OFC represent value signals if they are not required for behavior? One potential answer is that OFC computes and represents inferred values in all situations, and that these signals may bias choices at any point (Ballesta et al., 2020). However, if direct experience is available, these signals are typically indistinguishable from, and redundant with, cached values represented elsewhere in the brain, such that disruption of OFC does not affect observed behavior. In contrast, because no other brain area computes model-based values, OFC becomes important for behavior when outcomes must be inferred. This proposal would explain why animals and humans with compromised OFC function are capable of making choices, but that these choices reflect previously learned values even if they are no longer valid (Gallagher et al., 1999; Izquierdo et al., 2004; West et al., 2011; Rudebeck et al., 2013; Murray et al., 2015; Gardner et al., 2017; Reber et al., 2017; Gardner et al., 2018; Parkes et al., 2018; Howard et al., 2020).

In line with our previous work showing neural correlates of inferred outcomes in OFC (Wang et al., 2020), the current findings suggest that OFC networks are directly involved in stepping through the cue-cue and cue-reward associations when inferring outcomes at the time of decision making. However, alternative explanations have been proposed that do not require inference at this time point. For instance, cue A could be reactivated at the time of conditioning, such that it also acquires model-free value, just like cue B. Several studies have provided correlative evidence for such mediated learning processes in areas of the medial prefrontal cortex and temporal lobe (Shohamy and Wagner, 2008; Wimmer and Shohamy, 2012; Zeithamova et al., 2012; Kurth-Nelson et al., 2015). The strongest evidence against such mediated learning comes from reports that cue A does not support conditioned reinforcement (Sharpe et al., 2017), and that responding to this cue is sensitive to devaluation (Hart et al., 2020), the gold standards for assessing model-free and model-based value, respectively. Moreover, pharmacological inactivation of the OFC in the probe test selectively disrupts responding to cue A without affecting responding to cue B (Jones et al., 2012). If responding to both A and B were based on the same neural mechanisms involving model-free values, then presumably the two would not be differentially affected by OFC inactivation in the final probe test in this earlier experiment or, indeed, in the current study.

However, it is important to keep in mind that behavior can be driven by several independent mechanisms and that inference-based behavior may occur in parallel with support from additional mechanisms such as mediated learning (Schlichting and Preston, 2015), which may recruit hippocampus (Shohamy and Wagner, 2008; Wimmer and Shohamy, 2012; Kurth-Nelson et al., 2015) and perirhinal cortex (Wong et al., 2019). Nevertheless, the susceptibility of inference-based responding to OFC-targeted cTBS indicates that at least some amount of behavior in our task is based on real-time model-based computations. In this regard, it is important to note that whereas OFC-targeted cTBS reduced subjects’ ability to make inference based decisions, it did not fully abolish this function. This could be related to the fact that we only applied unilateral stimulation, and thus the contralateral OFC network may have remained unimpaired. Alternatively, the remaining performance could be driven by mediated learning processes mentioned above, dependent on areas not impacted by our OFC-targeted manipulation.

A limitation of our study is that we did not measure rsfMRI directly after TMS. Therefore, although stimulation sites were selected to have maximal connectivity with the central-lateral OFC, and we have previously shown that an identical OFC-targeted protocol disrupts activity in the lateral OFC network (Howard et al., 2020), we ae not able to confirm that this was the case in our current sample. It is therefore possible that local effects of our stimulation on LPFC drove the observed effects. However, we think this is unlikely for the following reasons. First, our TMS protocol was identical to our previous study in which we did not observe any effects on LPFC activity (Howard et al., 2020). Second, our results parallel previous findings with pharmacological inactivation of OFC in animals (Jones et al., 2012). Third, although medial PFC networks have been implicated in inference processes (Zeithamova et al., 2012; Schlichting et al., 2015; Schlichting and Preston, 2015), we are not aware of similar findings related to LPFC. However, cTBS could have affected reliability signals in LPFC that have been shown to correlate with the arbitration of behavioral control between model-based and model-free processes (Lee et al., 2014).

An additional limitation is our sham condition, which involved stimulation at 5% RMT. This is noticeably different from stimulation at 80% RMT in terms of auditory and somatosensory stimulation. Thus, these unintended peripheral effects of TMS could have driven the observed behavioral effects, rather than the neural changes induced by cTBS. We believe this is unlikely for two reasons. First, effects of cTBS were specific to inference-based behavior, and no differences were found for behavioral responses based on direct experience or memory for cue-cue associations. It is difficult to conceive why peripheral effects of the TMS would have highly disparate effects on two almost identical behaviors that only differ in their requirement for inference. Second, our previous study utilizing OFC-targeted TMS involved an additional control condition that was matched for somatosensory stimulation (Howard et al., 2020). Despite comparable peripheral effects, behavioral and neural effects in this control condition differed significantly from active cTBS but were similar to the 5% sham condition. We therefore think it is unlikely that our results were driven by unintended non-neuronal effects of cTBS.

In summary, our results support the idea that human OFC networks are necessary for inference-based behavior, whereas they are not critical to support decision making when direct experience is available. Deficits in decision making are a hallmark of many neuropsychiatric disorders, including substance use disorder (SUD) (Volkow and Fowler, 2000; Franklin et al., 2002; Goldstein et al., 2007; Zilverstand et al., 2018), obsessive compulsive disorder (OCD) (Menzies et al., 2008; Gillan et al., 2011; Nakao et al., 2014). Our findings may offer a conceptual framework for understanding how OFC dysfunction may disrupt behavior in these conditions. For instance, an impaired ability to imagine unobservable states may reinforce checking behaviors in OCD, and a failure to infer the consequences of long-term drug-use may bias drug-taking decisions in SUD. It would be important to develop OFC-targeted TMS protocols that enhance rather than disrupt OFC network activity with the goal to develop novel treatments that target specific behavioral dysfunctions in these disorders.

## Acknowledgments

The authors thank Rachel Reynolds, Devyn E. Smith, and Kelly Vogel for help with data collection, and Molly Hermiller for technical support related to TMS. This work was supported by grants from the National Institute on Deafness and other Communication Disorders (NIDCD, R01DC015426 to T.K.), the National Institute on Drug Abuse (NIDA, R03DA040668 to T.K.), and the Intramural Research Program at NIDA (ZIA-DA000587 to G.S.). The opinions expressed in this article are the authors’ own and do not reflect the view of the NIH/DHHS. The funders had no role in study design, data collection and analysis, decision to publish, or preparation of the manuscript.

## Author Contributions

F.W., G.S., J.L.V, and T.K. designed the experiment. F.W. and J.D.H. collected the data. F.W. analyzed the data. F.W., G.S. and T.K. interpreted the results and wrote the manuscript.

